# An atlas of human viruses provides new insights into diversity and tissue tropism of human viruses

**DOI:** 10.1101/2021.11.01.466721

**Authors:** Sifan Ye, Congyu Lu, Ye Qiu, Heping Zheng, Xingyi Ge, Aiping Wu, Zanxian Xia, Taijiao Jiang, Haizhen Zhu, Yousong Peng

## Abstract

Viruses continue to threaten human health. Yet, the complete viral species carried by humans and their infection characteristics have not been fully revealed. This study curated an atlas of human viruses from public databases and literatures, and built the Human Virus Database (HVD) available at http://computationalbiology.cn/humanVirusBase/#/. The HVD contains 1,131 virus species of 54 viral families which were more than twice the number of the human-infecting virus species reported in previous studies. These viruses were identified in human samples including 68 human tissues, the excreta and body fluid. The viral diversity in humans was age-dependent with a peak in the infant and a valley in the teenager. The tissue range of viruses was found to be associated with several factors including the viral group (DNA, RNA or reverse-transcribing viruses), enveloped or not, viral genome length and GC content, viral receptors and the virus-interacting proteins. Finally, the tissue range of DNA viruses was predicted using a random-forest algorithm with a medium performance. Overall, the study not only provides a valuable resource for further studies of human viruses, but also deepens our understanding towards the diversity and tissue tropism of human viruses.

## Introduction

Viruses are the most abundant and diverse biological entities on the earth (Jian et al., 2021). About 10^31^ viral particles are existing in all habitats of the world at any given time (Paez-Espino et al., 2016). Viruses have extensive host range, diverse virion morphology, and various genome types. The genomes of viruses can be DNA or RNA; linear or circular; and with length ranging from thousands to million bases. The shape of viruses can be predominantly grouped into filaments and spheres. In addition, viruses can infect all types of organisms, from animals and plants to bacteria and archaea (Dimmock et al., 2016; Liang and Bushman, 2021).

Among all viruses, human viruses have been intensively studied, especially those that cause disease and death. The most notorious human viruses include the influenza viruses (Paules and Subbarao, 2017), Ebola virus (Malvy et al., 2019), coronaviruses (Cui et al., 2019), and so on. For example, the 1918 Spanish Flu caused by the influenza A(H1N1) virus was reported to cause at least 50 million human death globally (Tumpey et al., 2005); the current COVID-19 pandemic caused by SARS-CoV-2 has resulted in more than 200 million infections and 4 million deaths as of August 6^th^, 2021 (WHO, 2021). Besides these viruses, more than 300 virus species of 26 viral families have been reported to infect humans (Mihara et al., 2016; Mollentze et al., 2021; Rodrigues et al., 2017), including both the human and zoonotic viruses (for clarity, both of them were considered as human viruses in the study). However, the list of human viruses is far from complete. The Global Virome Project estimated that there were 631,000-827,000 viruses in birds and mammals with the potential of human infection (Carroll et al., 2018). Therefore, it is in great need to identify more human viruses.

The rapid development of next-generation-sequencing (NGS) technology has enabled identification of novel viruses at an unprecedented rate compared to traditional methods of virus identification based on virus isolation (Cantalupo and Pipas, 2019). Both the DNA and RNA sequencing can be used to identify viruses. For example, Shi et al. identified 1,445 novel RNA viruses from over 220 invertebrate species based on the RNA-seq technology, which redefined the invertebrate RNA virosphere (Shi et al., 2016). Especially, lots of viruses have been identified in humans using NGS. For example, Zhou et al. identified the SARS-CoV-2 from the COVID-19 patients by the RNA-seq (Zhou et al., 2020), and Moustafa et al. identified 19 viruses by whole-genome sequencing of blood from 8,240 individuals (Moustafa et al., 2017).

The increasingly accumulated NGS data in the public databases provide a rich resource for identifying human viruses. Simon et al. built the MetaMap resource which contained numerous microbial and viral reads identified from more than 17,000 human disease-related RNA-seq samples deposited in the NCBI SRA database (Simon et al., 2018), while Kumata et al. detected 39 viral species from 8,991 RNA-seq data obtained from 51 somatic tissues of 547 healthy individuals (Kumata et al., 2020). A recent work by Gregory et al. constructed the human Gut Virome Database (GVD) which included 33,242 viral populations (only 2% were eukaryotic viruses) identified by assembly of 2,697 gut metagenomes from 32 studies (Gregory et al., 2020). Although much progress has been made on human viruses, there is still a lack of an integrated database for human viruses.

Human viruses can infect nearly all tissues of the human body (Liang and Bushman, 2021). The factors determining the tissue tropism of viruses have been investigated in previous studies, which can be grouped into viral and host factors. In terms of viral factors, the viral receptor-binding protein (RBP) contributes most to the viral tissue tropism (Hulswit et al., 2016; Maginnis, 2018). For example, influenza viruses infect tissues or cells with either α2,3-linked or α2,6-linked sialylated glycans, which is determined by the structure of hemagglutinin (the RBP of influenza viruses) (Tzarum et al., 2015). Besides, other viral proteins can also have an influence on the viral tissue tropism. For example, the flavivirus NS1 protein can affect the virus tissue tropism by disruption of endothelial glycocalyx components (Puerta-Guardo et al., 2019). The host factors determining the tissue tropism of viruses have been summarized in McFadden’s study (McFadden et al., 2009), and can be grouped into three kinds: 1) the virus receptor which is responsible for viral entry into host cells. For example, the NTCP, the receptor of Hepatitis B virus (HBV), is mainly expressed in liver and determines the strong hepatotropism of HBV (Li, 2015); 2) antiviral genes which limit the virus invasion, such as the interferons; 3) host factors indispensable to viral replication, transcription and other steps in viral life cycle. However, only a few viruses have been comprehensively studied for their tissue tropism.

This study curated an atlas of human viruses from public databases and literature, and built the Human Virus Database (HVD) for storing them; then, the viral diversity in human tissues and the dynamics of viral diversity by age were analyzed; then, the tissue range of viruses and the factors contributing to the viral tissue range were investigated; finally, a model of predicting the tissue range of DNA viruses was built. The study not only provides a resource for exploring the human viruses, but also deepens our understanding towards the diversity and tissue tropism of human viruses.

## Materials and methods

### 1 Data source of human viruses

The human viruses were curated from four sources: 1) the NCBI GenBank database. All viral sequences were firstly downloaded with the GenBank format from the NCBI GenBank database on June 26^th^, 2021. Those which were isolated from human samples were kept by manual curation. The tissues from which the viruses were identified were extracted from the metadata if available. The viral sequences with less than 300 bp were removed. The viruses whose hosts are registered as invertebrates, vertebrates, and humans were defined as animal viruses. Only the animal viruses were kept for further analysis. This resulted in a total of 912 virus species which were isolated from humans based on the NCBI GenBank database; 2) the MetaMap database (Simon et al., 2018). The MetaMap database is an atlas of microbial and viral reads which were computationally extracted from the human disease-related RNA-seq data. The animal virus (see the definition mentioned above) which was detected in at least two projects and whose median expression abundance in multiple samples of the same project was no less than 1 RPM in at least one project was kept for further analysis. The meta data of viruses including the age, sex and the tissue of the samples from which the viruses were identified were extracted directly from the database; 3) the tissue level atlas of the healthy human virome from the Kumata’s study (Kumata et al., 2020). All 39 virus species identified in the study were collected except the Tomato spotted wilt tospovirus which was reported to infect both the invertebrates and the plants. 4) the human GVD (Gregory et al., 2020). The viral sequences in GVD were downloaded firstly; then, they were queried against the animal virus sequences by BLASTN. The viral sequences which had E-value ≦ 1E-5, identity ≧ 0.9, and coverage ≧ 0.8 to the best hits were kept, and they were considered to belong to the same viral species with the best hits; then, the viruses were further filtered by virus abundance. Only those which were detected in at least two projects and whose median raw abundance (defined as the average read depth in Gregory’s study) in multiple samples of the same project was no less than 3 in at least one project were kept.

All the viruses mentioned above were combined together and were organized by viral species and by human tissues. The tissues were grouped into eleven human systems. The following viruses were removed due to the possible contamination: viruses of the *Baculoviridae* family which are commonly used in the laboratory (Zapatka et al., 2020); the human endogenous retroviruses such as the Human endogenous retrovirus W, H and K; the animal retroviruses such as the Porcine type-C oncovirus, Murine leukemia virus and Abelson murine leukemia virus (Kearney et al., 2012; Moustafa et al., 2017); the Shamonda and Simbu orthobunyavirus which have some genomic sequences identical to human rRNAs and may be identified as false positives (Cantalupo et al., 2018); the Macaca mulatta polyomavirus 1 (SV40) which is commonly used in the plasmid (Cantalupo et al., 2018).

### 2 The biological features of viruses

The human viruses collected in this study were roughly grouped into the DNA, RNA and reverse-transcribing (RT) viruses according to the Baltimore classification system. Whether the virus is enveloped or not was determined based on the ViralZone database (Hulo et al., 2011). The genome sequences were obtained for 607 viruses from the NCBI RefSeq database on July 1^st^, 2021.

### 3 The receptor of human viruses

The receptor of 81 human viruses were obtained from the viralReceptor database (Zhang et al., 2019) on January 11^th^, 2021.

### 4 Protein-protein interactions between human and viruses

The protein-protein interactions (PPIs) between human and viruses were obtained from the Lasso’s study during which the high-confidence PPIs between human and 1,001 human-infecting viral strains were provided (Lasso et al., 2019). Only the high-confidence PPIs which have likelihood ratio (LR) values greater than 100 were used. The PPIs between human and multiple viral strains of the same viral species were combined together.

### 5 Immune-related genes

The immune-related genes in human were obtained from the database of InnateDB and ImmPort (Bhattacharya et al., 2018; Breuer et al., 2013) on April 26^th^, 2021.

### 6 The expression level of human genes in common human tissues

The expression level of human genes in 32 common human tissues were obtained from the Expression Atlas database (ID: E-MTAB-2836) (Papatheodorou et al., 2020) on Nov 26^th^, 2020.

### 7 Predicting the tissue range of DNA viruses with the random forest algorithm

The DNA viruses were classified into two groups based on the tissue range: one group of viruses infecting only one tissue and the other group of viruses infecting two or more tissues. The random forest (RF) algorithm was used to classify these two groups of viruses. The algorithm was achieved with the package of sklearn in Python (version 3.7) with default parameters. The leave-one-out tests by species, genus and family were used to evaluate the ability of the RF models in predicting tissue range for novel viruses. For example, in the leave-one-out test by genus, each viral genus was left out for testing and the remaining viral genera were used to built the RF model; then, the RF model was tested on the left-out genus. The AUC, accuracy, sensitivity and specificity were used to measure the performance of the RF model.

### 8 Statistical analysis

All the statistical analyses were conducted in R (version 3.2.5). The Wilcoxon rank-sum test was conducted by the function of *wilcox*.*test()* in R. The correlation coefficient was calculated by the function of *cor*.*test()* in R.

## Results

### 1 Virome diversity in humans

A total of 1,131 virus species were identified in humans. They were derived from four sources including the NCBI GenBank database, the MetaMap database, the Kumata’s study of identifying viruses from the healthy humans, and the human GVD (see Materials and Methods). Most of them (912/1,131) were derived from the NCBI GenBank database (Figure S1); about one-fourth of them (299/1,131) were derived from the MetaMap database. 93 virus species were derived from both the NCBI GenBank database and the MetaMap database (Figure S1).

The 1,131 human virus species included 433 DNA, 656 RNA and 42 RT viruses according to the Baltimore classification system (Table S1). They could be further classified into 54 families of which *Picornaviridae* was the largest family and contained 314 virus species. The top 10 largest viral families (Figure S2) contained 65% of all virus species.

By isolation source, a total of 701 and 296 virus species were isolated from human tissues or cells, and the human excreta or body fluid, respectively. Among them, 92 virus species were isolated from both sources. Besides, a total of 226 virus species of 32 viral families were identified in humans without detailed sources. Analysis of the viruses identified in human tissues showed that although some viral families like the *Picornaviridae, Papillomaviridae, Anelloviridae* and *Herpesviridae* were abundant in multiple tissues, which is consistent with previous studies, most viral families were only abundant in one or a few tissues (Figure 1). For example, the *Hantaviridae* was only abundant in the blood. On the other hand, most tissues contained viruses from multiple viral families, although the virus number varied much across tissues. The cardiovascular system which includes tissues of blood, heart, bone marrow and blood vessel contained the largest number of virus species. The blood contained the largest number of viral species (381) among all tissues, and the other tissues (heart, bone marrow, blood vessel) also contained more than 50 virus species from multiple viral families such as the *Herpesviridae* and *Retroviridae*. The respiratory system which includes the pharynx, lung, nose, trachea bronchi, larynx and chest wall contained the second largest number of viral species among all systems. For example, the lung contained 114 viral species from 29 viral families, especially the *Herpesviridae*. The digestive system which includes 14 tissues such as the liver, colon, ileum, esophagus, and so on, contained the third largest number of viral species among all systems. For example, the liver contained 102 viral species from 26 viral families, especially the *Herpesviridae* and *Retroviridae*. Interestingly, several important internal tissues including the brain, prostate gland, uterus, kidney and bone contained more than 50 viral species from multiple viral families. For example, the brain which is protected by the blood-brain barrier and was supposed to be sterile contained 139 viruses from more than 10 viral families.

**Figure 1.**
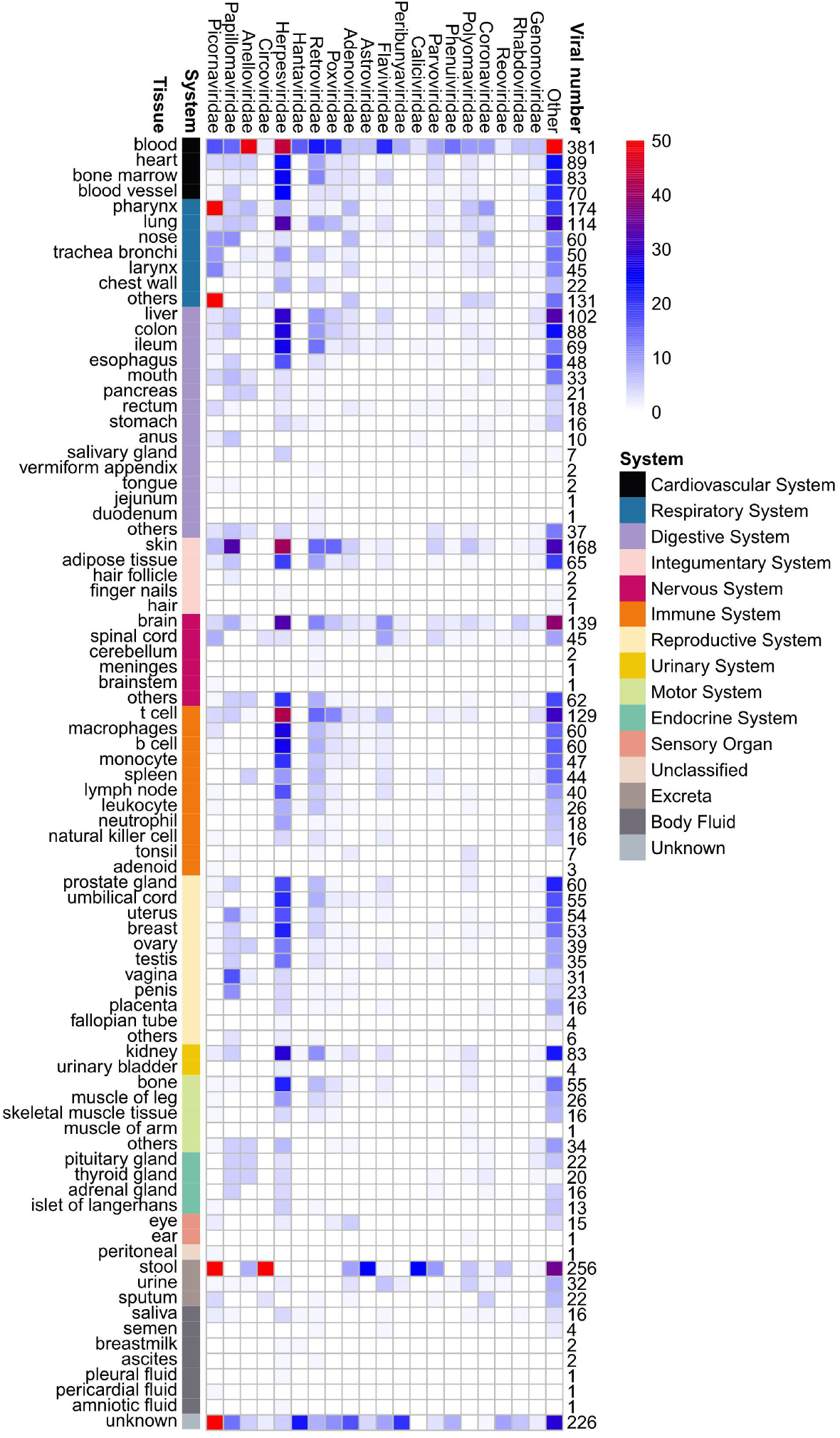
Viruses identified in humans tissues, excreta and body fluid. The tissues were shown in the left side which were grouped by systems. “Others” referred to other tissues or cells of the system combined together. The top 20 largest viral families were shown in the top of the figure. For each viral family, the number of viral species detected in a tissue was colored according to the legend in the top-right. The numbers in the right side showed the total number of viral species detected in a tissue.

In addition to the viruses identified in human tissues, a total of 296 virus species were identified in the human excreta including the stool, urine, and sputum, and in the body fluid including the saliva, semen, breast milk, and so on. The stool contained 256 virus species from more than 10 viral families, especially the *Picornaviridae* and *Circoviridae*. Besides, the urine and sputum also contained 32 and 22 viruses, respectively. However, the body fluid contained only a few viruses except the saliva.

### 2 Temporal dynamics of virome diversity throughout the life

The dynamics of viromes by sex and age were analyzed. No significant difference was observed between the number of viral species per sample in the male and female (Figure S3). Besides, the composition of viral families detected in the samples of males and females was also similar (Figure S4). Therefore, the samples of males and females were combined together in analysis of the temporal dynamics of viromes in individuals of different life stages. Although the overall composition of viral families were similar in samples of different life stages (Figure S5), the number of viral species per sample varied a lot throughout the life. As shown in Figure 2, the median number of viral species per sample was 17 in the infant (0-3 years old); then, it decreased to 4 in teenagers (3-18 years old); in adults (18-65 years old), it increased to 15; finally, it decreased a little in elders (>65 years old). This suggested that the virome dynamics pattern was age-dependent which was similar to that observed for the virome diversity in the human gut.

**Figure 2.**
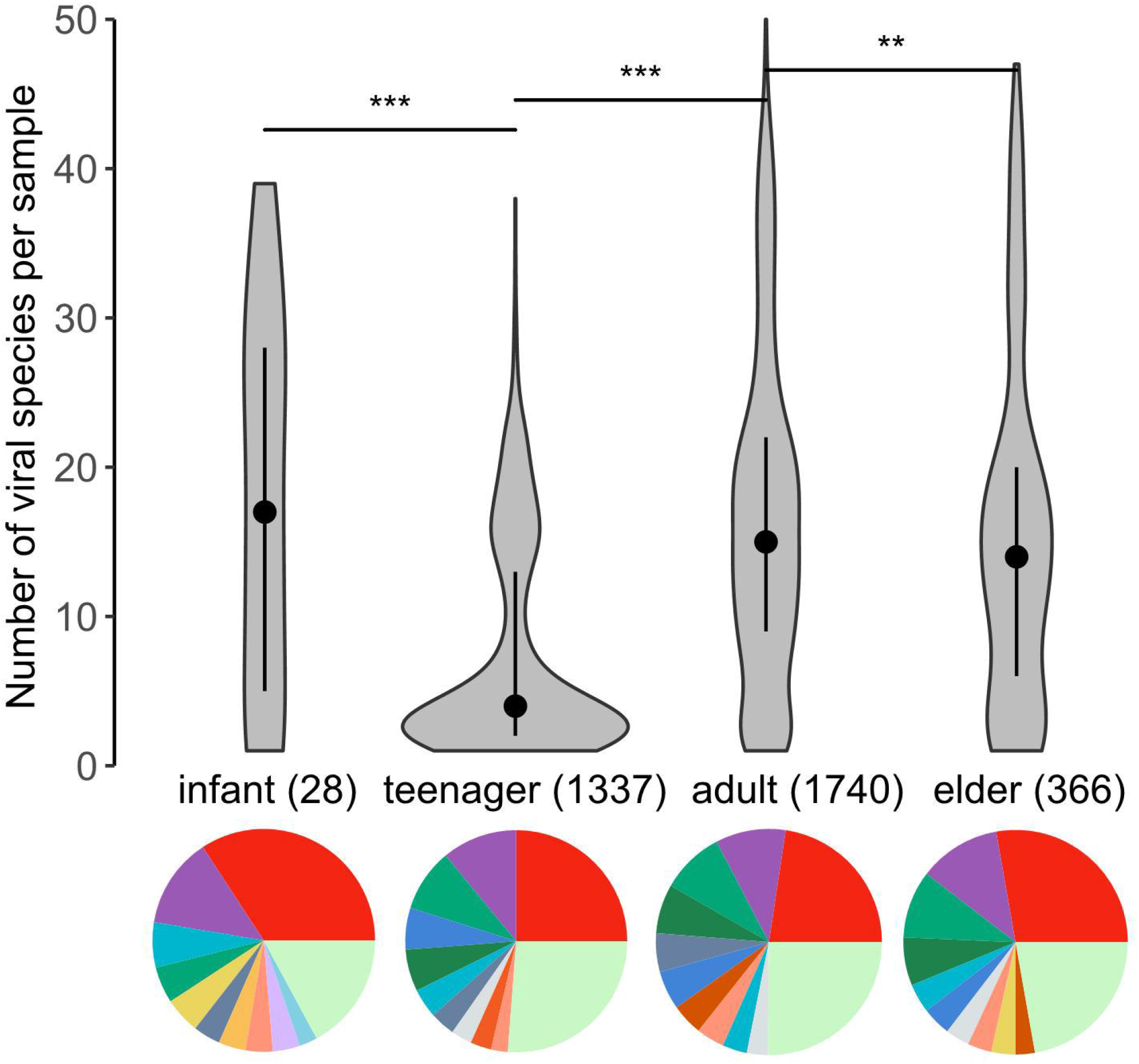
The virome dynamics in different life stages of humans. Only the viruses derived from the MetaMap database were used as the metadata of viruses are of high-quality in the database. The number in the brackets referred to the number of samples used in the life stage. The pie charts in the bottom showed the composition of viral families detected in samples of different life stages. Please see Figure S5 for the larger version of the pie charts. **, p-value < 0.01; ***, p-value < 0.001.

### 3 The tissue range of viruses

The tissue range of human viruses was analyzed. As shown in Figure 3A, the tissue range varied much among the 701 virus species which were identified in human tissues or cells. More than half of virus species infected only one tissue and 72.8% of virus species infected less than five tissues, while 90 virus species were observed to infect 10 or more tissues, such as the Ictalurid herpesvirus 1 and the Cotesia congregata bracovirus. The factors which may influence the tissue range of viruses were analyzed as following.

**Figure 3.**
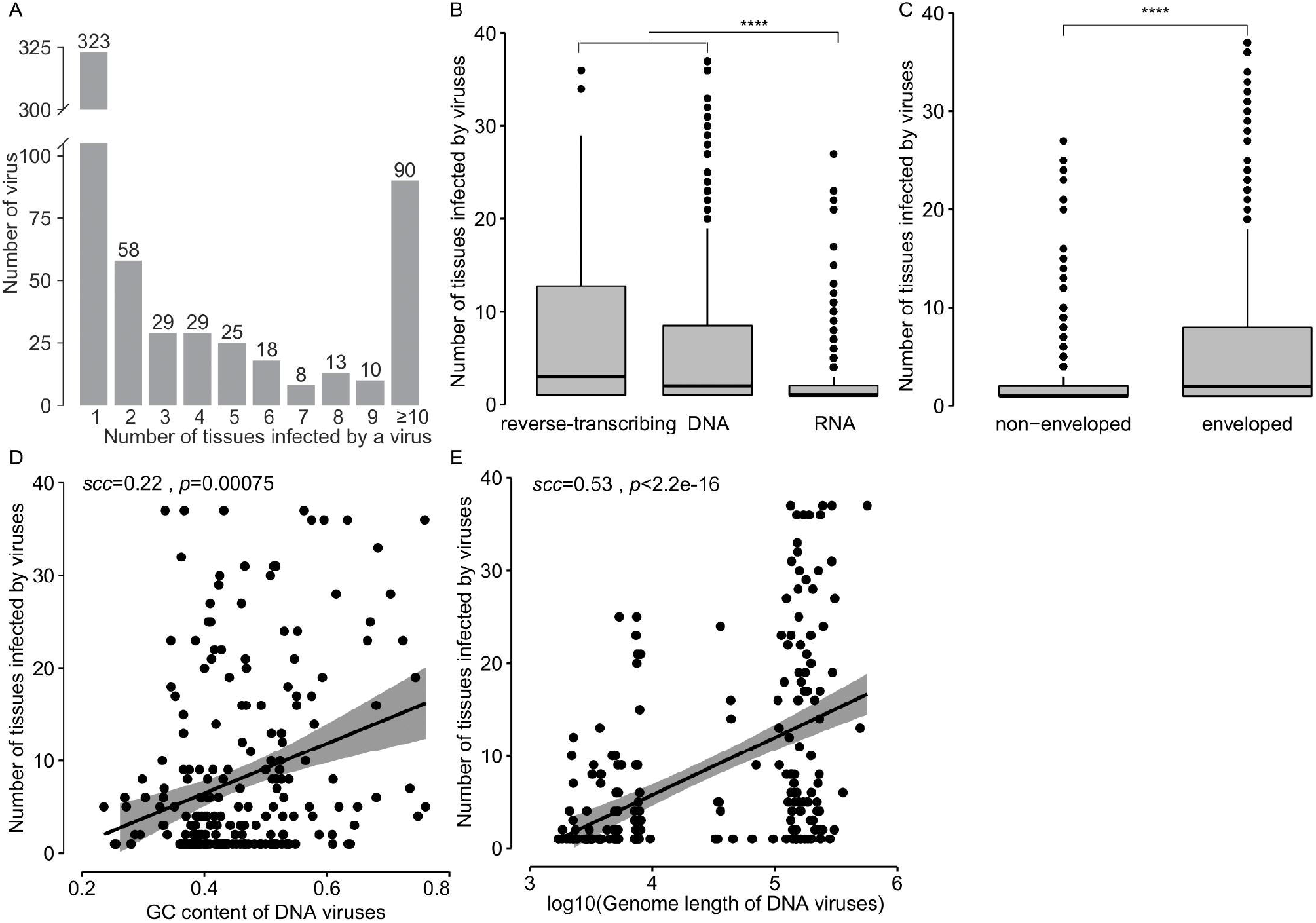
The biological factors influencing the tissue range of human viruses. (A) The distribution of the number of human tissues infected by viruses. (B) Comparison of the number of tissues infected by DNA, RNA and RT viruses; (C) Comparison of the number of tissues infected by enveloped and non-enveloped viruses; (D) Correlation between the GC content and tissue range of DNA viruses; (E) Correlations between the genome length and tissue range of DNA viruses. The SCCs and the related p-values were shown in the top-left of the figures. The lines in (D) and (E) referred to the linear least square regression lines, and the gray regions referred to the 95% confidence intervals. ****, p-value < 0.0001.

#### 1) Biological factors

The tissue range of different groups of viruses was analyzed firstly. The RT viruses infect a range of 1-36 tissues, with the first quartile (Q1), median, third quartile (Q3) being 1, 3, 12.8, respectively; the DNA viruses infected a range of 1-37 tissues, with the Q1, median, Q3 being 1, 2, 8.5, respectively; the RNA viruses infected a range of 1-27 tissues, with the Q1, median, Q3 being 1, 1, 2, respectively. The tissue range of both the RT and DNA viruses was significantly larger than that of RNA viruses (p-value < 0.0001 in the Wilcoxon rank-sum test) (Figure 3B).

Then, the tissue range of enveloped or non-enveloped viruses was analyzed. The enveloped viruses infected a range of 1-37 tissues, with the Q1, median, Q3 being 1, 2, 8, respectively; the non-enveloped viruses infected a range of 1-27 tissues, with the Q1, median, Q3 being 1, 1, 2, respectively. The tissue range of the enveloped viruses was significantly larger than that of the non-enveloped viruses (p-value < 0.0001 in the Wilcoxon rank-sum test) (Figure 3C).

Then, the genome features were analyzed for viruses of different tissue range. The GC content of viral genomic sequences was observed to have a weak positive correlation with the tissue range of viruses, with the Spearman correlation coefficient (SCC) of 0.17 (p-value < 0.001) (Figure S6). When analyzing the relationship between the virus tissue range and GC content by viral group, a weak correlation was observed for DNA viruses with a SCC of 0.24 (p-value < 0.001) (Figure 3D), while no correlations were observed for both the RNA and RT viruses (Figure S6). Further analysis of the relationship between the genome size and the tissue range of viruses showed a weak positive correlation with a SCC of 0.33 (p-value < 0.001) (Figure S7A). When analyzing the relationship between the genome size and the number of tissues infected by viruses by viral group showed a middle positive correlation for DNA viruses with a SCC of 0.53 (Figure 3E), while no correlations for RNA and RT viruses (Figure S7B&C).

#### 2) Viral receptors

Viral receptors are key to viral infection of hosts. Previous studies have shown that the virus may use more than one receptors (Wang et al., 2021). Among the viruses used here, 48 viruses had more than one receptor, and 33 viruses used one receptor. We hypothesized that the more receptors used by a virus, the larger the tissue range is. As expected, a significant positive correlation (SCC=0.49, p-value=0.02) was observed between the number of viral receptors and the number of tissues infected by DNA viruses (Figure 4A), while no correlations were observed for RNA and RT viruses (Figure S8). We next investigated whether the expression level of viral receptors had an influence on the tissue range of viruses. The expression levels of viral receptors in 32 common human tissues were analyzed. As expected, the expression levels of viral receptors in the infected tissues were a little higher than those in the non-infected tissues (p-value < 0.05) (Figure S9). A weak positive correlation (SCC=0.30, p-value=0.05) was observed between the number of tissues infected by viruses and the median expression levels of viral receptors used by the virus (Figure S10A). However, when analyzing the associations in terms of DNA, RNA or RT viruses, no significant correlations were observed (Figure 4B and Figure S10B&C).

**Figure 4.**
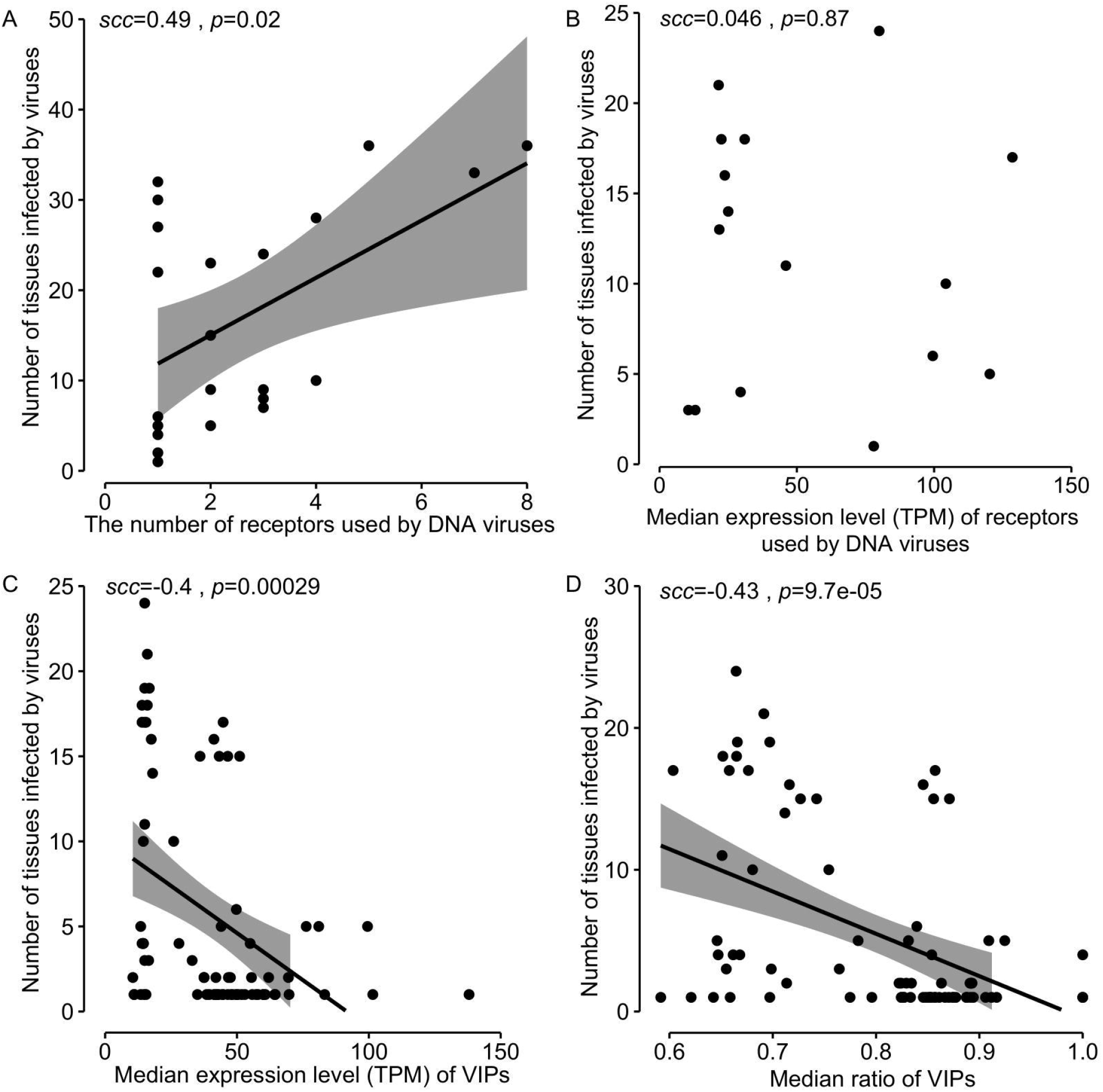
The influence of viral receptors and virus-interacting proteins (defined as VIPs) on the tissue range of DNA viruses. (A)-(D) referred to the correlations between the number of tissues infected by viruses and the number of receptors used by the virus (A), the expression level of viral receptors (B) and VIPs (C), the ratio of VIPs(D). The SCCs and the related p-values were shown in the top-left of the figures. The lines in (A), (C) and (D) referred to the linear least square regression lines, and the gray regions referred to the 95% confidence intervals. Only 32 common human tissues were used in the analysis shown in figures B-D as the gene expression levels were only available in these tissues.

#### 3) Virus-interacting proteins

Besides the viral receptors, there are lots of human proteins interacting with viral proteins which may also have an influence on the tissue range of viruses. Thus, we obtained the PPIs between human and viruses from the Lasso’s study during which the high-confidence PPIs between human and 1,001 human-infecting viral strains were predicted. A total of 5,449 virus-interacting proteins (defined as VIPs) were obtained for 132 virus species, including 78 DNA viruses, 51 RNA viruses and 3 RT viruses. We hypothesized that the expression levels of VIPs may have an influence on the tissue range of viruses. The correlations between the number of tissues infected by viruses and the median expression level of VIPs in 32 common tissues were analyzed for DNA and RNA viruses. Unexpectedly, a negative correlation was observed (SCC =-0.40, p-value=0.00029) for DNA viruses (Figure 4C), while no significant correlation was observed for RNA viruses (Figure S11A). Considering that the VIPs with low expressions may contribute little to viral infection, we calculated the ratio of VIPs which had median or high expressions in each tissue (defined as ≧ 4 TPM which is the median expression level of all genes in 32 human tissues), and analyzed the correlation between the number of tissues infected by viruses and the median ratio of VIPs with median or high expressions in 32 common tissues. Again, a negative correlation was observed with a SCC of −0.43 for DNA viruses (Figure 4D), while no significant correlation was observed for RNA viruses (Figure S11B).

The high expression of immune-related genes may inhibit viral infection. We then investigated whether the immune-related genes contributed to the negative correlations observed above. 972 of the total 5,449 VIPs were involved in the innnate or (and) adaptive immunity in humans. Thus, the VIPs were classified as the immune-related VIPs and non-immune-related VIPs. The relationships between the number of tissues infected by viruses and the expression levels (or ratio) of immune-related or non-immune-related VIPs in 32 common tissues were analyzed. Similar results were obtained as above: for DNA viruses, the negative correlations were observed for both the immune-related and non-immune-related VIPs (Figure S12), while for RNA viruses no significant correlations were observed for both kinds of VIPs (Figure S13).

### 4 Prediction of tissue range of DNA viruses

Since the DNA viruses showed significant correlations between the number of tissues infected by viruses and multiple factors, we further investigated the prediction of the tissue range for DNA viruses. The DNA viruses were separated into two groups: one group of viruses infecting one tissue, while the other group of viruses infecting more than one tissue. The random forest (RF) algorithm was used to classify these two groups of DNA viruses based on five features including the GC content, genome length, enveloped or not, the median expression level of VIPs, and the ratio of VIPs with median or high expressions in 32 common human tissues. The features related to viral receptors were not used in the modeling because only 81 viruses had receptors identified. A total of 105 viruses with all five features which included 37 viruses infecting one tissue and 68 viruses infecting two or more tissues were used in the modeling. The leave-one-out tests by species, genus and family were used to evaluate the ability of the RF model in predicting tissue range for novel viruses (see Materials and Methods). As shown in Table 1, the RF model had both the AUCs and accuracies greater than 0.7 at both the species and genus level in the leave-one-out tests, while at the family level the performance of the RF model decreased much, with the AUC and accuracy being 0.66 and 0.69, respectively.

**Table 1.** The performance of the RF model in predicting the tissue range of DNA viruses in the leave-one-out tests by species, genus and family.

| Leave-one-out test | AUC | Accuracy | Sensitivity | Specificity |
| --- | --- | --- | --- | --- |
| By species | 0.79 | 0.72 | 0.81 | 0.57 |
| By genus | 0.74 | 0.70 | 0.81 | 0.49 |
| By family | 0.66 | 0.69 | 0.86 | 0.35 |

### 5 Overview of the HVD

A database named Human Virus Database (HVD) was created to store and organize the human viruses. It is freely available to the public at http://computationalbiology.cn/humanVirusBase/#/. The HVD mainly includes Home, Browse, Search, Statistic, Download and Tutorial pages (Figure 5).

**Figure 5.**
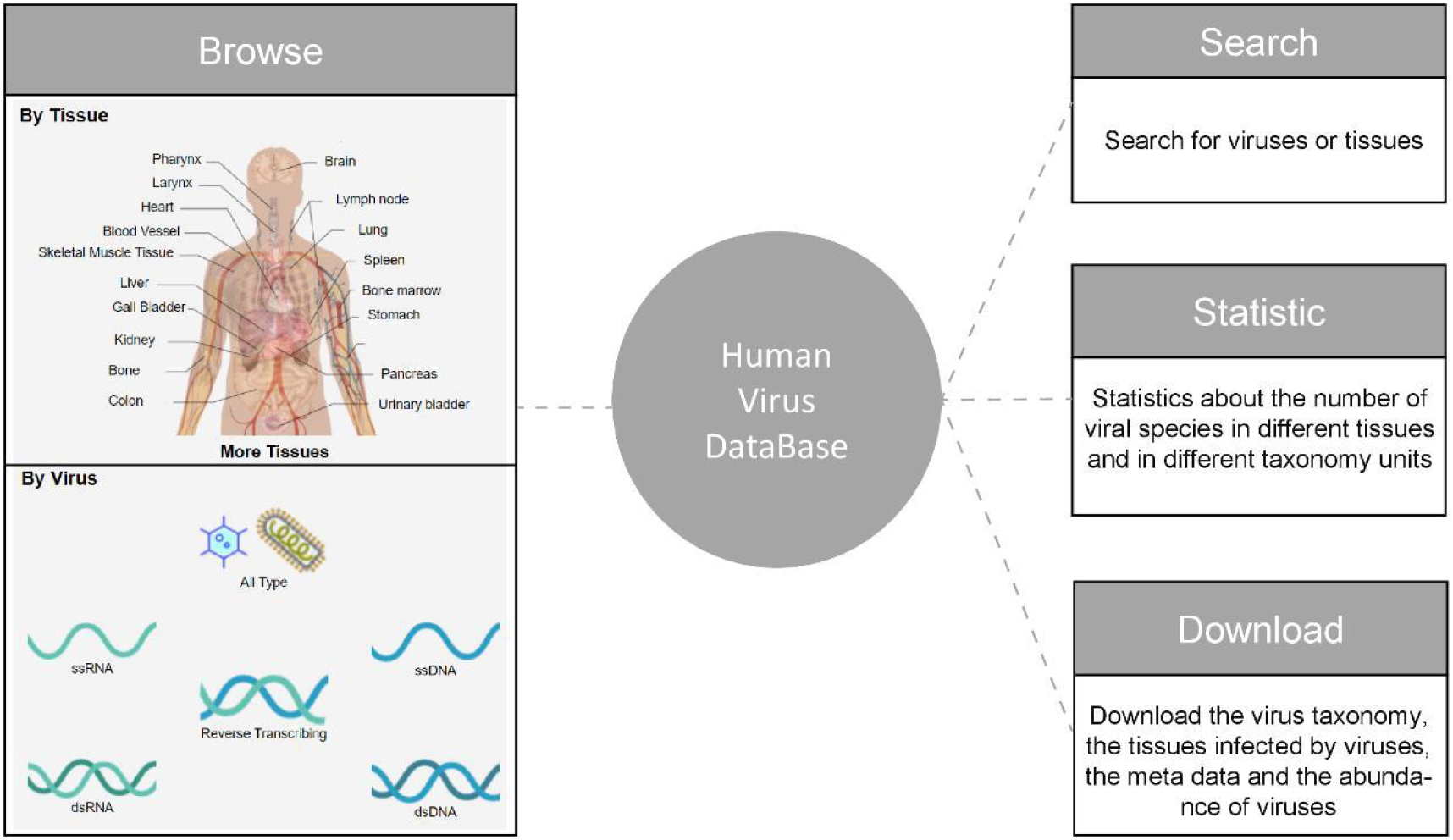
The structure of the HVD.

#### Browse

The page displayed the human virus by tissue or by viral taxonomy. When browsing the virus by tissue, the viruses identified in the tissue were shown in a table; the taxonomy distribution of these viruses by family was shown in a pie chart; and the abundance of these viruses (if given) were shown in box plots. When browsing the virus by viral taxonomy, the viruses were firstly organized by the Baltimore classification system and then by viral family. For each virus, the following information was provided in a table: the tissues infected by the virus were organized by human systems; the metadata such as age, sex and location related to the virus isolation (if given); the evidence of the virus identification in the tissues; the median abundance of the virus (if given). Besides, the tissue distribution of the virus was shown in a pie chart and the abundances of the virus in tissues were shown in a box plot.

#### Search

Users can search for viruses or tissues by name.

#### Statistic

This page displayed a summary statistics about the number of virus species identified in each tissue in 11 systems, and the number of viral species in each viral taxonomy unit.

#### Download

All the information stored in the database, including the virus taxonomy, the tissues infected by viruses, the meta data and the abundance of viruses, were freely available for downloading by the tissue or by viral family.

## Discussion

This study presented an up-to-date atlas of viruses detected in human bodies. More than 1,000 virus species were identified in humans which was much larger than that of the human-infecting virus species reported in previous studies (Mihara et al., 2016; Mollentze et al., 2021; Rodrigues et al., 2017). This suggested that much more viruses are supposed to infect humans than expected. Actually, the Global Virome Project estimated that there were 631,000-827,000 viruses with the potential of human infection. The viruses identified in the study should be put in priority due to their potential of human infection.

Many factors determine the tissue range of viruses. We found the RT and DNA viruses infected a wider range of tissues than the RNA viruses; the enveloped viruses infected more tissues than the non-enveloped viruses. Interestingly, there was a median to strong correlation between the number of tissues infected by the virus and the genome size for DNA viruses. DNA viruses generally have much larger genomes than RNA viruses, thus encoding lots of proteins. For example, the Epstein-Barr virus has a genome of 171,823 bp and encodes more than 80 proteins (Dolan et al., 2006). The more proteins the virus encodes, the less the virus relies on host cells. Besides, a lot of viral proteins have been reported to counteract the immune system of the host cells. Our analysis showed that 96.4% of proteins in DNA viruses interacted with the immune-related VIPs (data not shown). Thus, the viruses with larger genomes could encode more proteins, which may help the viruses defeat the host immune system.

The receptor was reported to be a key factor for determining the tissue tropism of viruses. For DNA viruses, a median positive correlation was observed between the number of tissues infected by viruses and the number of receptors used by the virus, while for RNA viruses and RT viruses no significant correlations were observed. This was possibly because the DNA viruses with large size need more receptors for entry into host cells, or because the viral receptors identified are incomplete yet. Weak or no significant correlations were observed between the number of tissues infected by viruses and the expression of viral receptors. This suggests that the expression level of viral receptors may have little influence on the tissue range of viruses considering that viral receptors have much higher expressions than other cell membrane proteins in common human tissues (Zhang et al., 2019).

The virus-host interactions were also reported to be important factors determining the tissue tropism of viruses. Interestingly, we observed significant negative correlations between the number of infected tissues and the expression level or the ratio of VIPs for DNA viruses. The immune-related VIPs could inhibit the viral infections and are supposed to have negative correlations with viral tissue range (McFadden et al., 2009). However, when the analysis was conducted for the non-immune-related VIPs, the negative correlation was still observed. This suggests that most VIPs are suspected to inhibit the DNA virus infection.

There are some limitations in this study. Firstly, the human viruses identified in this study may be far from complete when compared to the estimate by the Global Virome Project. Much more human viruses would be discovered in the future with the NGS-based method. Nevertheless, this study provided an up-to-date atlas of human viruses which were stored and organized in the user-friendly HVD. It would help much in further studies of human viruses. Secondly, most viruses obtained in this study were identified based on the NGS method. They need further experimental validations. Thirdly, the HVD only contained animal viruses in humans and ignored the phages which are abundant in humans.

In conclusion, this study built the HVD which included 1,131 virus species identified in humans. Large viral diversity was observed in multiple tissues of humans, and the virome diversity was age-dependent with peak in the infant and valley in the teenager. The tissue range of viruses was found to be associated with several factors including the viral group, enveloped or not, viral genome length, viral receptors and VIPs. Overall, this study not only provided a valuable resource for further studies of human viruses and for early warnings of newly emerging viruses, but also deepened our understanding towards the diversity and tissue tropism of human viruses.

### Key Points

- An atlas of human viruses was curated and stored in the Human Virus Database.
- The viral diversity in humans was age-dependent with a peak in the infant and a valley in the teenager.
- The tissue range of viruses was associated with viral group, enveloped or not, viral genome length and GC content, viral receptors and the virus-interacting proteins.

## Acknowledgements

This work was supported by the National Key Plan for Scientific Research and Development of China (2016YFD0500300), Hunan Provincial Natural Science Foundation of China (2020JJ3006), the Key Research and Development Program of Hunan Province (2020SK2054) and the National Natural Science Foundation of China (82072293)

## Compliance with Ethical Standards

### Animal and Human Rights Statement

This article does not contain any studies with human or animal subjects performed by the author.

### Biographical Note

Sifan Ye is a master students in the College of Biology, Hunan University.

Congyu Lu is a PhD candidate in the College of Biology, Hunan University.

Ye Qiu and Yousong Peng are associate professors in the College of Biology, Hunan University.

Heping Zheng, Xingyi Ge, Haizhen Zhu are professors in the College of Biology, Hunan University.

Aiping Wu and Taijiao Jiang are professors in Suzhou Institute of Systems Medicine.

Zanxian Xia is a professor in Central South University

