## Additional file for "An atlas of human viruses provides new insights into diversity and tissue tropism of human viruses"

### Supplementary Figures

**Figure S1** Source of viruses used in this study.

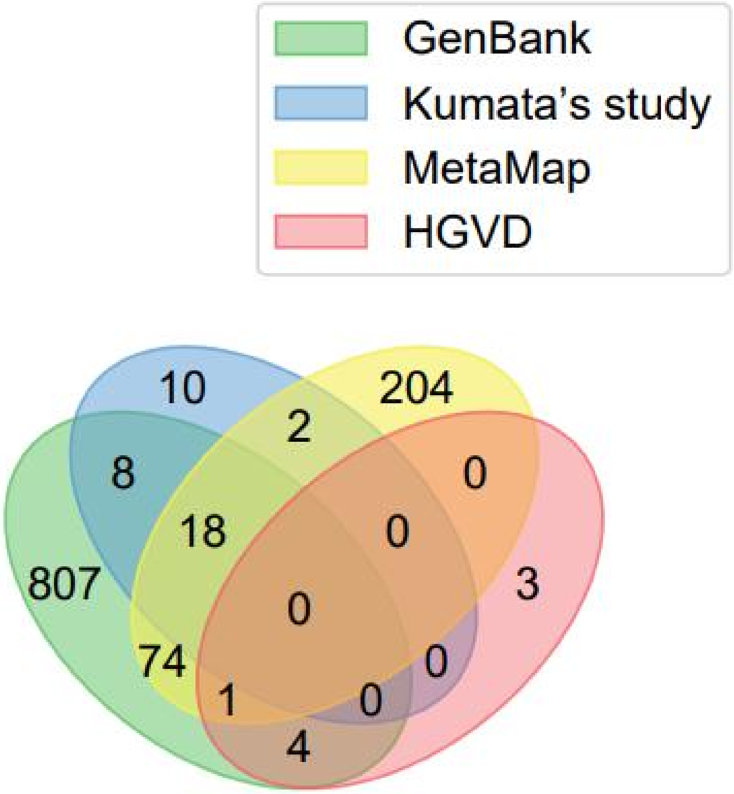

**Figure S2** Viral family composition of viruses used in this study.

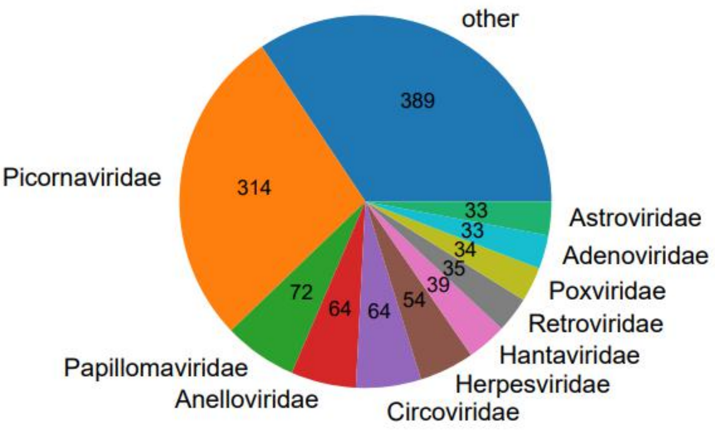


**Figure S3** The number of viral species per sample in males and females. Ns, no significant difference.

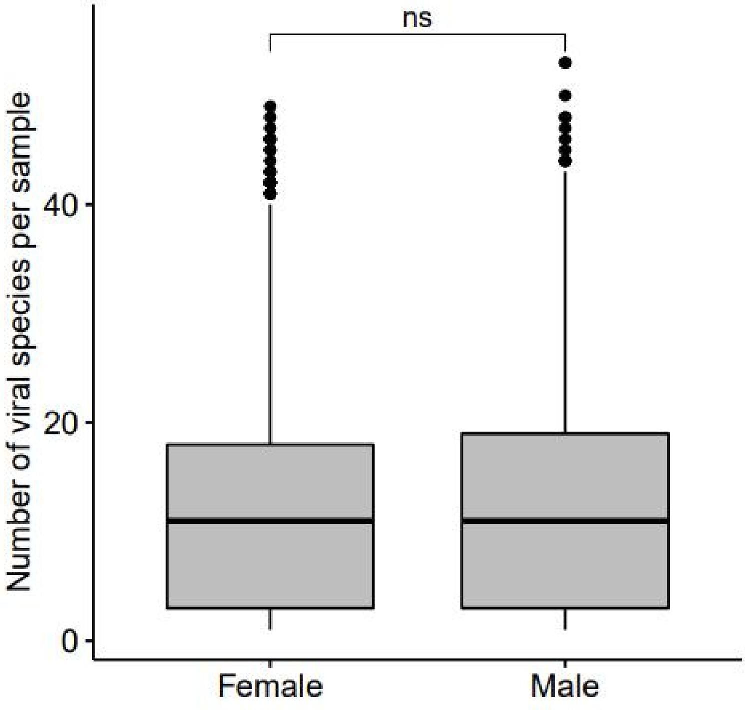

**Figure S4** The composition of viral families detected in samples of males and females.

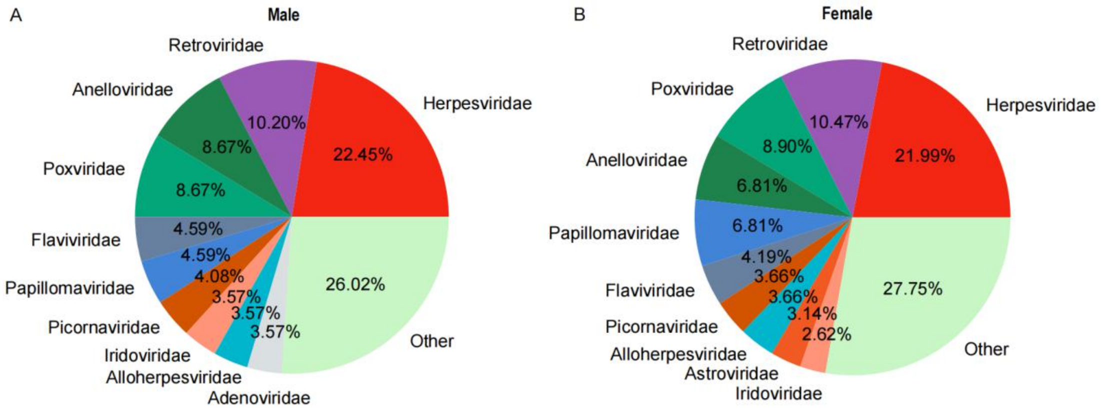


**Figure S5** The composition of viral families detected in samples of different life stages.

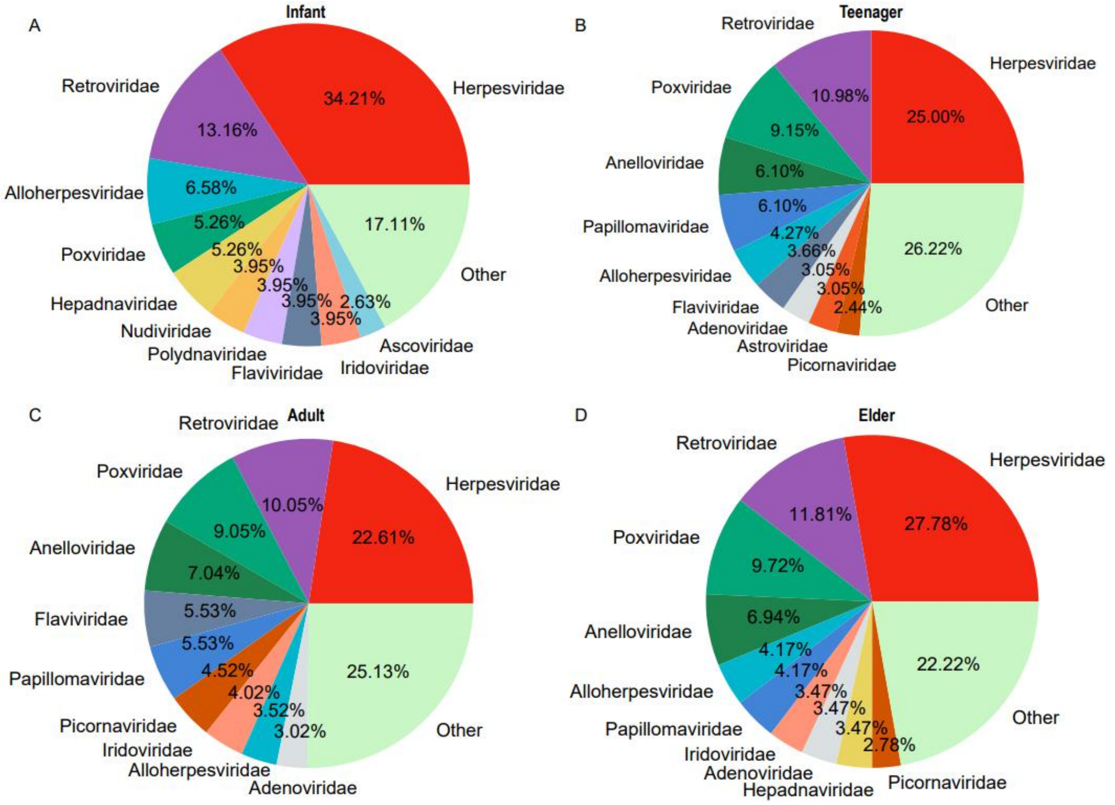

**Figure S6** The relationship between the GC content of viral genomic sequences and the number of tissues infected by all viruses (A), RNA viruses (B) and reverse-transcribing viruses (C).

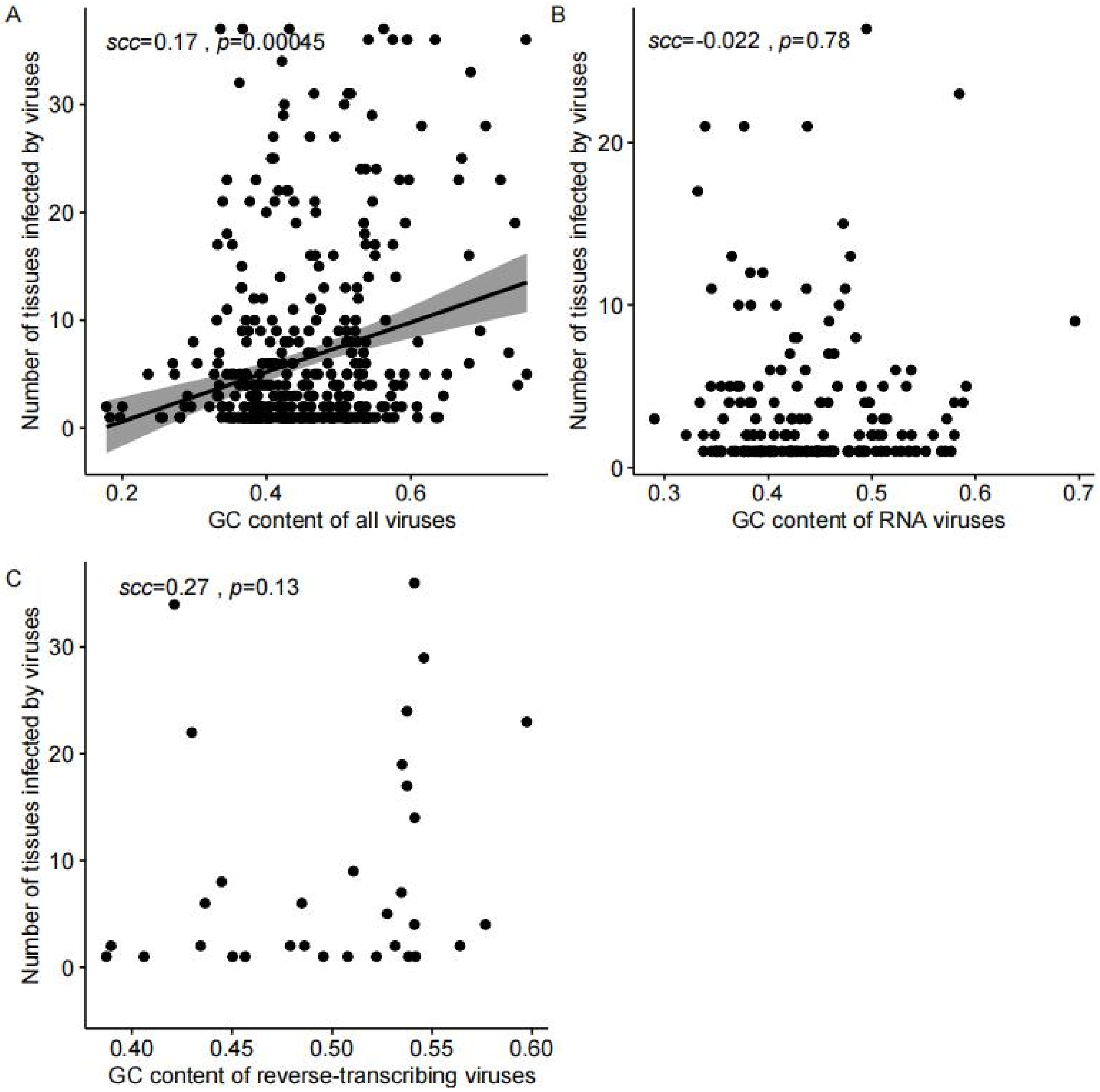

**Figure S7** The relationship between the genome length and the number of tissues infected by all viruses (A), RNA viruses (B) and reverse-transcribing viruses (C).

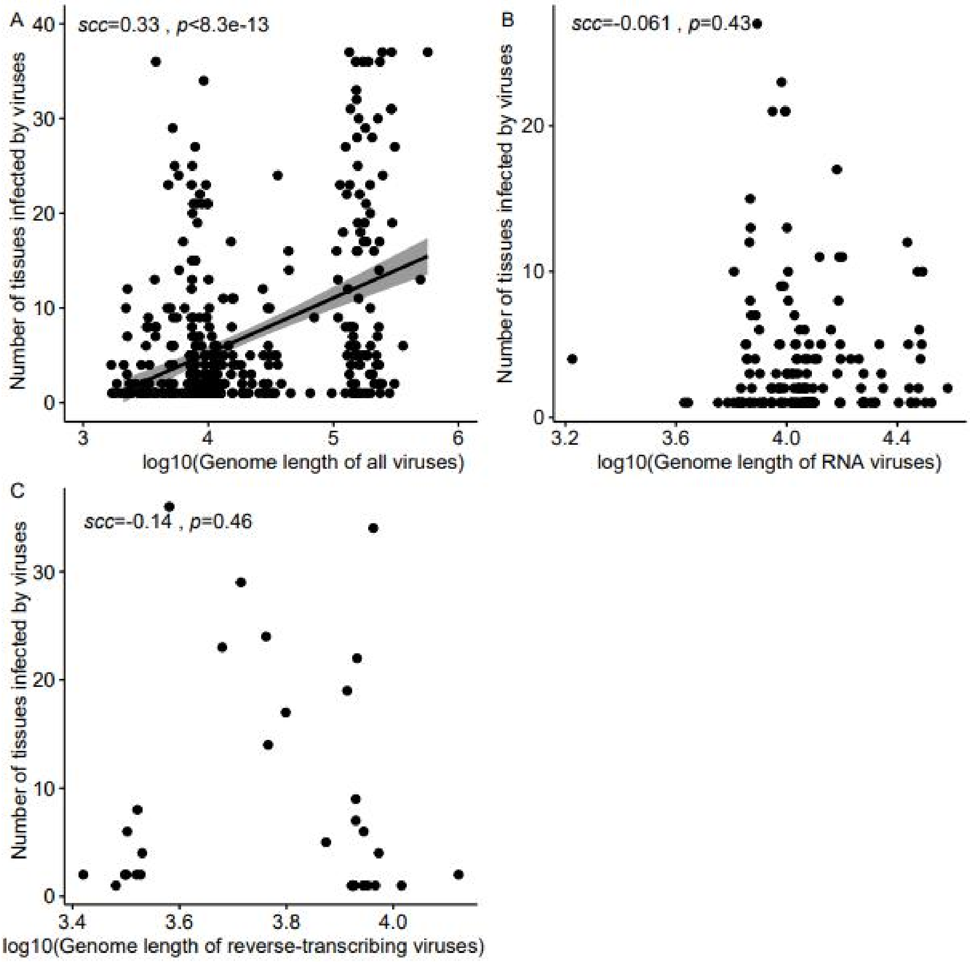

**Figure S8** The relationship between the number of viral receptors and the number of tissues infected by all viruses (A), RNA viruses (B) and reverse-transcribing viruses (C).

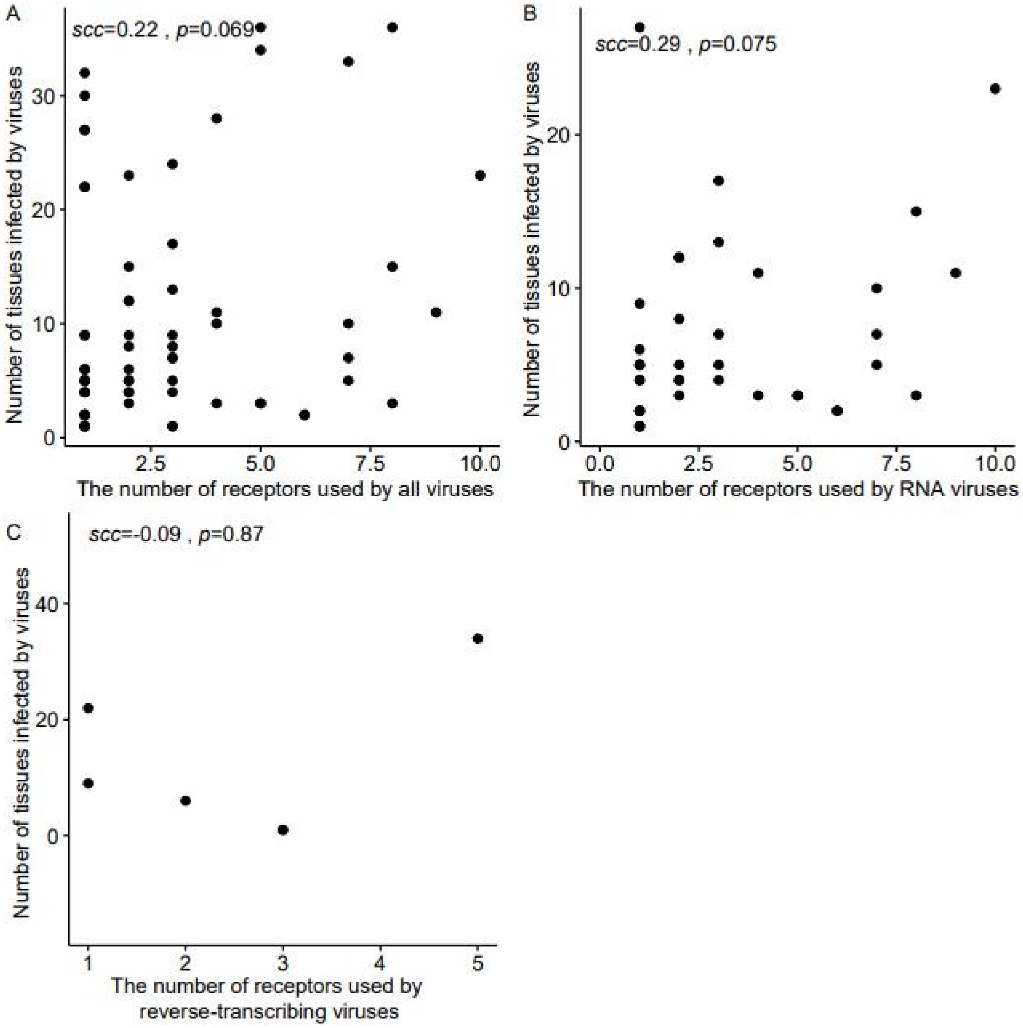

**Figure S9** Comparison of the expression levels of viral receptors in the infected tissues and non-infected tissues. *, p-value < 0.05.

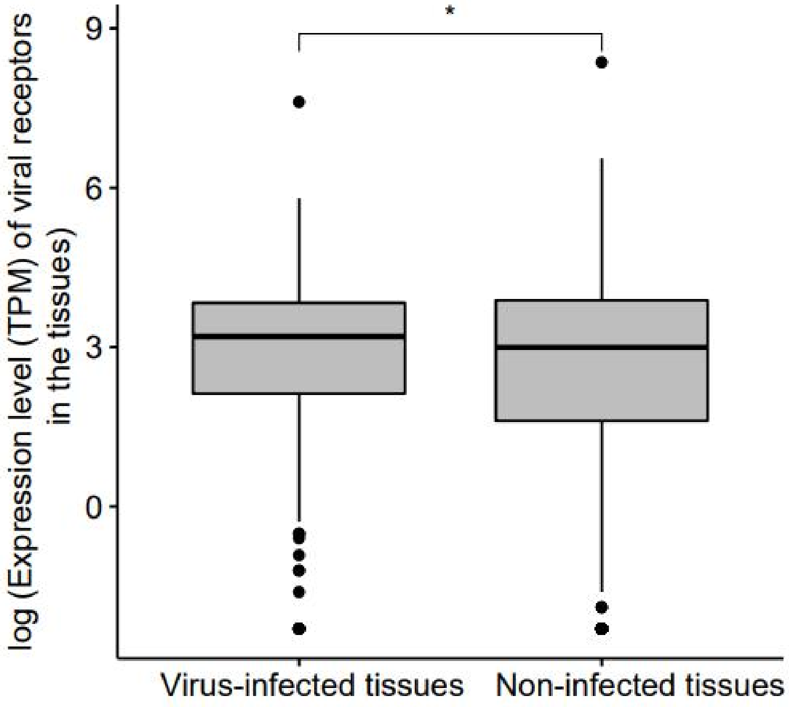

**Figure S10**. The relationship between the number of tissues infected by viruses and the median expression levels of viral receptors used by all viruses (A), RNA viruses (B) and reverse-transcribing viruses (C).

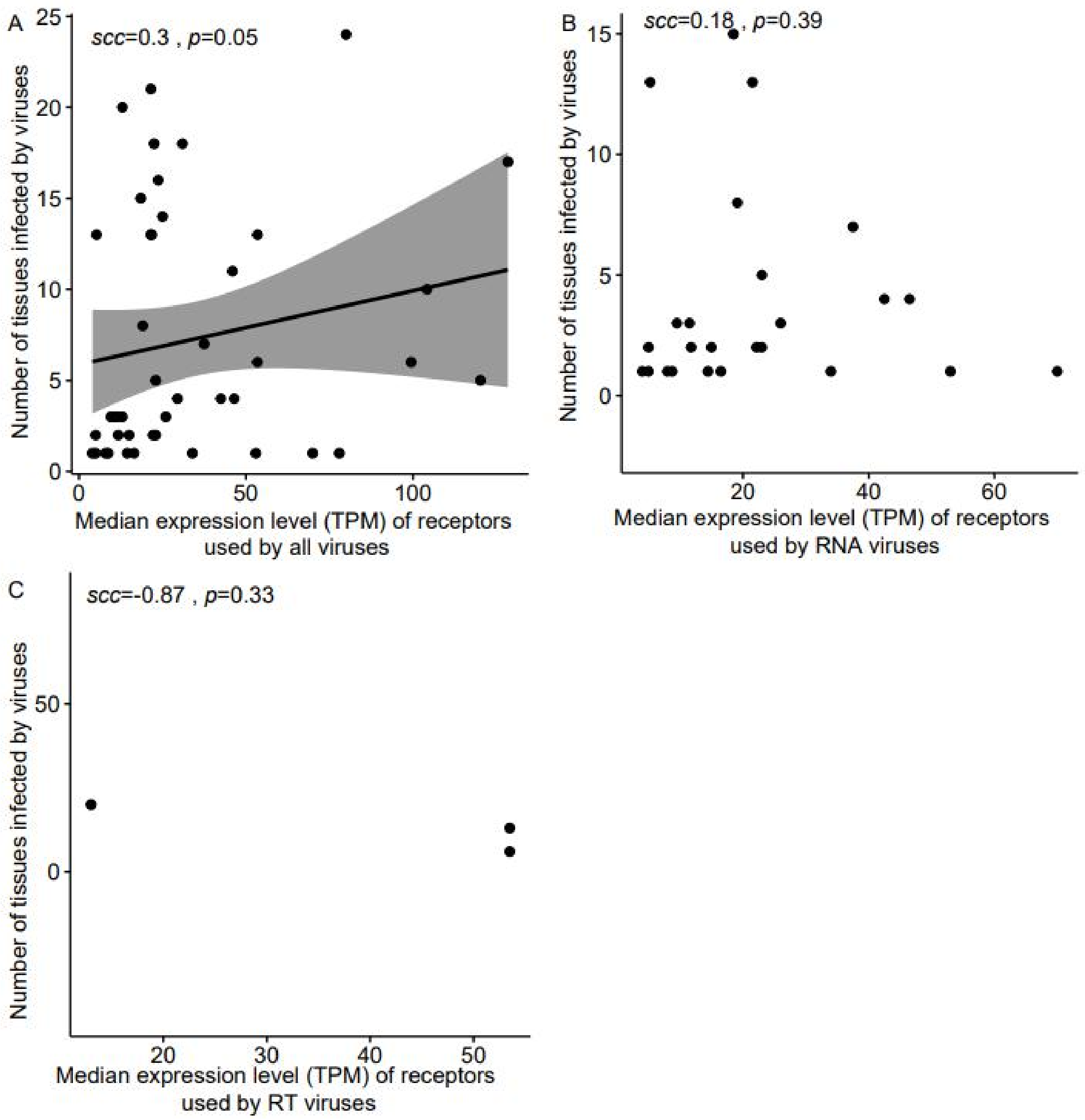

**Figure S11** The relationship between the number of tissues infected by RNA viruses and the median expressions level of VIPs (A), and the median ratio of VIPs (B) in 32 common tissues.

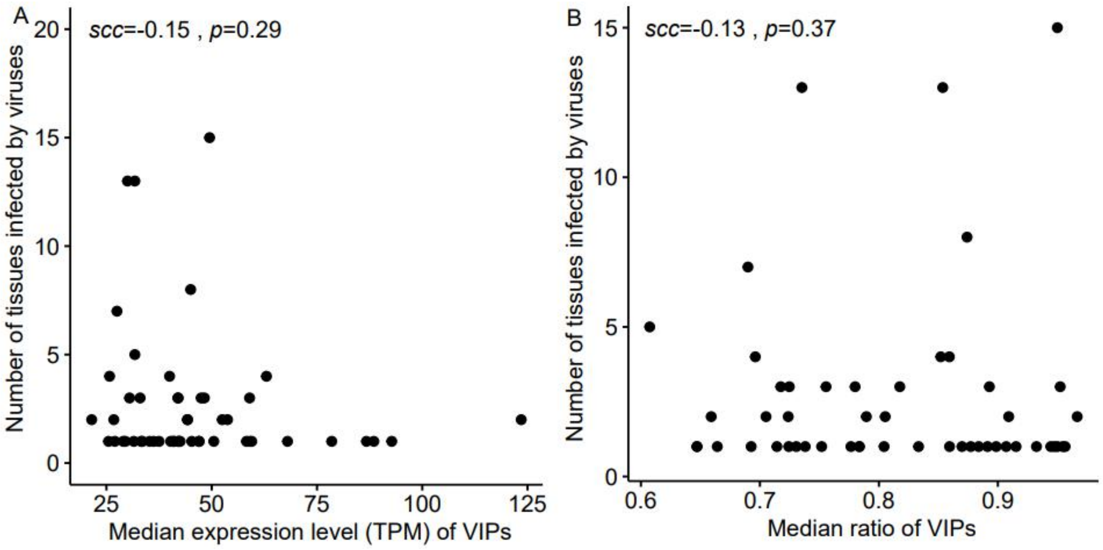

**Figure S12** The relationship between the number of tissues infected by DNA viruses and the median expressions level of immune-related VIPs (A) and non-immune-related VIPs (B), the median ratio of immune-related VIPs (C) and non-immune-related VIPs (D) in 32 common tissues.

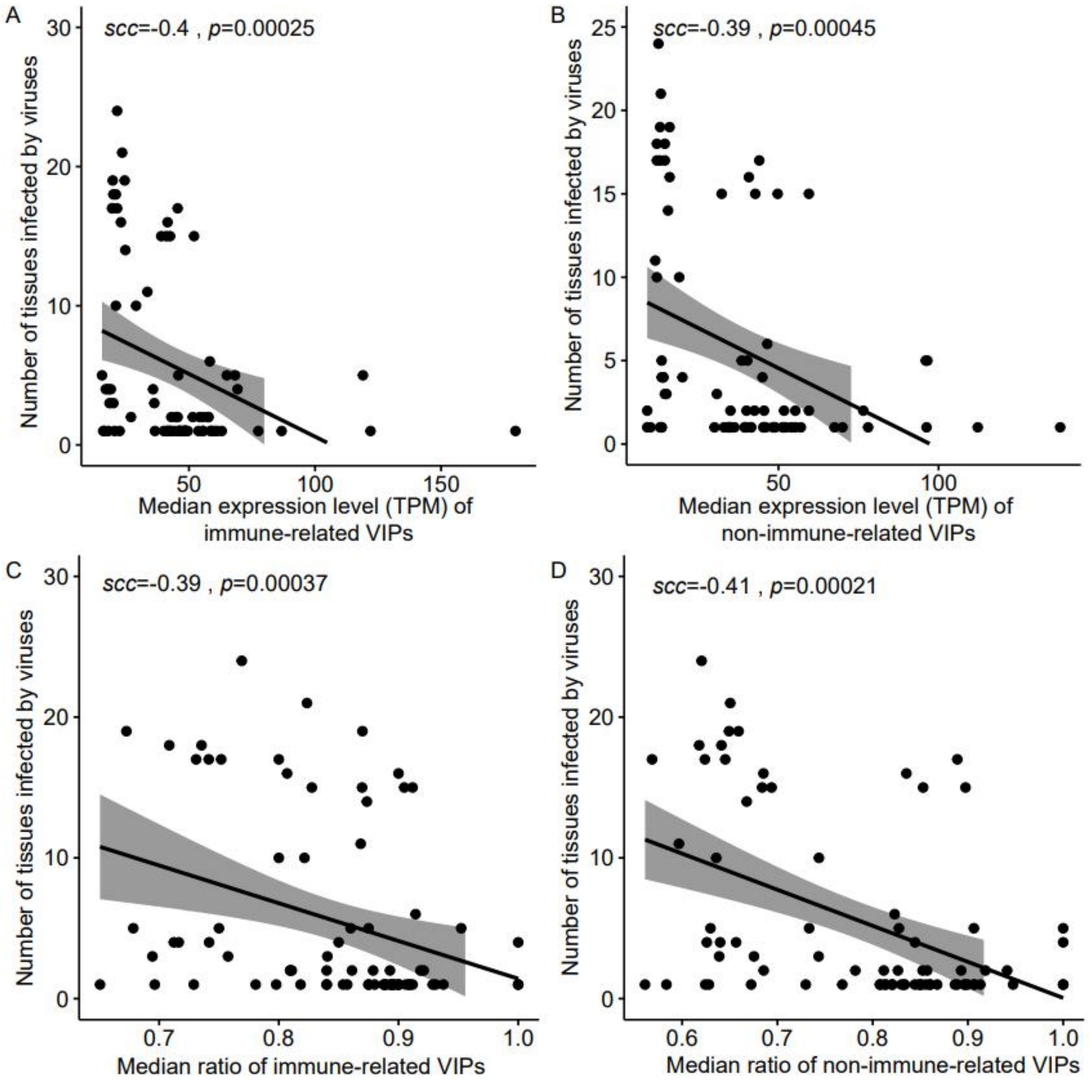

**Figure S13** The relationship between the number of tissues infected by RNA viruses and the median expressions level of immune-related VIPs (A) and non-immune-related VIPs (B), the median ratio of immune-related VIPs (C) and non-immune-related VIPs (D) in 32 common tissues.

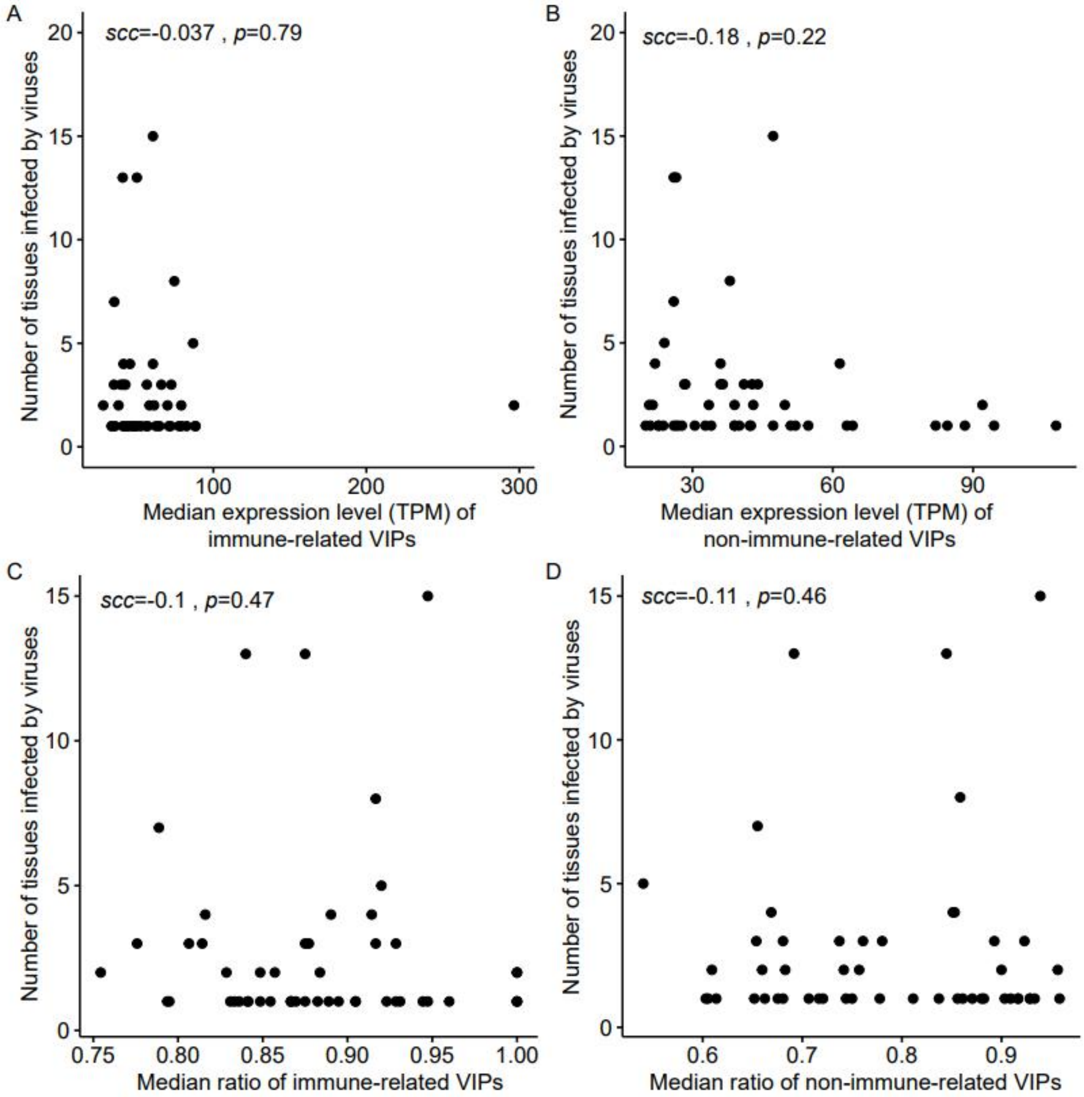
